# Biliary microbiota and bile acids composition in cholelithiasis

**DOI:** 10.1101/469825

**Authors:** Vyacheslav A. Petrov, María A. Fernández-Peralbo, Rico Derks, Elena M. Knyazeva, Nikolay V. Merzlikin, Alexey E. Sazonov, Oleg A. Mayboroda, Irina V. Saltykova

## Abstract

**Background:** A functional interplay between BAs and microbial composition in gut is a well-documented phenomenon. In bile this phenomenon is far less studied and with this report we describe the interactions between the BAs and microbiota in this complex biological matrix.

**Methodology:** Thirty-seven gallstone disease patients of which twenty-one with *Opisthorchis felineus* infection were enrolled in the study. The bile samples were obtained during laparoscopic cholecystectomy for gallstone disease operative treatment. Common bile acids composition were measured by LC-MS/MS using a column in reverse phase. For all patients gallbladder microbiota was previously analyzed with 16S rRNA gene sequencing on Illumina MiSeq platform. The associations between bile acids composition and microbiota were analysed.

**Principal findings:** Bile acids signature and *O. felineus* infection status exerts influence on beta-diversity of bile microbial community. Direct correlations were found between taurocholic acid, taurochenodeoxycholic acid concentrations and alpha-diversity of bile microbiota. Taurocholic acid and taurochenodeoxycholic acid both shows positive associations with the presence of Chitinophagaceae family, *Microbacterium* and *Lutibacterium* genera and *Prevotella intermedia*. Also direct associations were identified for taurocholic acid concentration and the presence of Actinomycetales and Bacteroidales orders, *Lautropia* genus, *Jeotgalicoccus psychrophilus* and *Haemophilus parainfluenzae* as well as for taurochenodeoxycholic acid and Acetobacteraceae family and Sphingomonas genus. There were no differences in bile acids concentrations between O. *felineus* infected and non-infected patients.

**Conclusions/Significance:** Associations between diversity, taxonomic profile of bile microbiota and bile acids levels were evidenced in patients with cholelithiasis. Increase of taurochenodeoxycholic acid and taurocholic acid concentration correlates with bile microbiota alpha-diversity and appearance of opportunistic pathogens. Alteration of bile acids signature could cause shifts in bile microbial community structure.

## Introduction

Liver bile ducts and gallbladder is one of the most uncharted biomes in human body due to invasiveness of its exploration. For long time bile of the healthy organisms was believed being sterile [1] but recent metagenomics studies identified a number of bacteria and archaea species which lives in intact bile ducts [2]. Since bile ducts do not have direct connections with the environment, the human bile flora most probably originated from upper digestive tract microbiome. It has, however, a lower taxonomic diversity consisting of *Enterobacteriaceae*, *Prevotellaceae*, *Streptococcaceae* and *Veillonellaceae* families [2].

A physiological status of the host is one of the key factors defining a composition of the bile microbiota. For instance, it has been shown that liver fluke infection results in elevation of taxonomic diversity in microbiota. Experimental infection of golden hamsters with *O. viverrini* results in alteration of taxonomic composition and increase of alpha-diversity in gut and bile microbiome [3]. Human-based study of gallbladder microbiota in *O. felineus* infection confirmed the fact of fluke-induced shifts in bile microbial community structure and introduction of taxons unseen in microbiota of non-infected, individuals [4]. Non-infectious pathologies strongly influence bile microbiota composition as well. It has been shown that primary sclerosing cholangitis leads to a reduction of microbial diversity, alteration of Pasteurellaceae, Staphylococcaceae, and Xanthomonadaceae abundances, while Streptococcus abundance shows a strong positive correlation with the disease severity and a number of the previous cholangiography examinations [5].

Moreover, a pathology driven shift in the microbiota diversity may lead to the alterations of the bile acids (BA) repertoire; in gut such interplay a system mictobiota-BA is well documented [6]. Alternatively, the BAs themselves can affect gut microbiota community directly (antimicrobial and progerminative actions) and indirectly via farnesoid X receptor activation [7].

In bile interplay between microbiota and BAs is much less studied. To date there is only a single report on cholangiolithiasis patients bile microbiota which provides evidence of an association between BAs levels and abundance of bacteria from *Bilophila* genus in the supraduodenal segment of common bile duct [8].

Here, to corroborate additional evidence of bile pathology-driving changes in gallbladder flora, we are aiming to explore the possible links between the most abundant BAs and microbiota on the background of cholelithiasis.

## Materials and methods

### Study population

The study was approved by Ethics Committee of the Siberian State Medical University. Thirty-seven participants of male and female gender, with age ranged from 40 to 61 and diagnosed with gallstone disease were enrolled in the study. Twenty-one of patients were diagnosed with *O. felineus* infection. The bile samples from all people were obtained during operative treatment of gallstone disease (laparoscopic cholecystectomy). During the surgical intervention 5–10 ml of gallbladder bile was aspirated under sterile conditions and immediately delivered to the laboratory. Two ml bile was clarified by centrifugation (10, 000 *g*, 10 min), the pellet was stored at -80°C for bile microbiota analysis, the supernatant was stored at -80°C for bile acid analysis.

### Bile acid analysis

LC-MS/MS analysis was applied for the quantification of the following ten of the most common bile acids (BAs) in a complex matrix such as the bile: cholic acid (CA), chenodeoxy cholic acid (CDCA), deoxycholic acid (DCA), ursodeoxycholic acid (UDCA), taurocholic acid (TCA), taurochenodeoxycholic acid (TCDCA), taurolithocholic acid (TLCA), glycocholic acid (GCA), glycochenodeoxycholic acid (GCDCA), glycodeoxycholic acid (GDCA). A detailed description of the analytical procedure is presented in the supplementary material.

### Bile microbiota analysis

Gallbladder bile samples microbiota for each of the patients were previously analyzed and described with 16S rRNA gene sequencing on Illumina MiSeq machine [4]. Raw 16S rRNA gene reads data are available on European Nucleotide Archive, accession number PRJEB12755, http://www.ebi.ac.uk/ena/data/view/PRJEB12755. Sequencing results were analyzed as described in Saltykova et al, 2016 [4]. Briefly, reads analysis was implemented in QIIME 1.9.0 [9] with the usage of the open reference OTU picking algorithm by the UCLUST method[10] and GreenGenes taxonomy v13.5 [11] as the reference base for taxonomic assignment. All OTUs present only in reagent control were subtracted from experimental samples to eliminate contamination. Samples with less than 200 sequences were removed from the study.

Alpha-diversity or microbial community taxonomic richness was calculated in QIIME using Chao1, Observed OTUs, Shannon and Simpson indices at depth of 200 sequences per sample. For further analysis all OTUs observed in less than 3 samples were excluded and microbial data was normalized with CSS algorithm [12]. Distances between samples in unweighted UniFrac metrics for estimation of pairwise dissimilarity between communities (beta-diversity) was calculated in QIIME.

### Statistical analysis

Statistical analysis was implemented in R 3.5.1 version [13]. To examine differences in BAs concentrations between infected and non-infected patients Mann-Whitney-Wilcoxon test was used. FDR-corrected Spearman rank correlation (psych package [14]) was used to define a linkage between taxonomic richness and BAs levels. Contribution of BAs concentrations to gallbladder flora beta-diversity estimated with permutational multivariate analysis of variance (algorithm adonis of vegan package [15]) with 9999 permutations in the model which includes distance matrix as outcome and transformed metabolites concentrations along with invasion status and gender as predictors. For this analysis BAs concentrations were transformed with non-metric multidimensional scaling (NMDS) in Euclidean metrics to one vector represents the value of first principle coordinate. NMDS was used for dimension reduction of microbial data to produce scatterplot for visualization of beta-diversity. All metabolites linked with microbiota diversity were enrolled in further analysis. Associations between microbial phylotypes and BAs levels were defined by linear regression in model with BAs concentrations as outcome, OTUs presence data as a predictor and age, gender, body mass index as covariates. In case of multiple hypotheses testing, p-values were corrected with FDR method. Visualization was made in ggplot2[16] and corrplot[17] packages.

## Results

### Gallbladder bile acids signature

Table 1 summaries the results of the LC-MS based quantification of the ten most abundant bile acids. The optimized and validated method was applied for the analysis of selected bile acids in the bile of 37 participants. Primary BAs and its conjugates have about 78% of abundance in gallbladder. The most abundant of measured BAs in human gallbladder with the concentrations more than 1000 ng/ml in all groups were GCA (31.9%), GCDCA (23.6%), GDCA (18.7%) and TCDCA (16.9%). Other measured BAs were amount to 9% of total concentration.

**Table 1.**
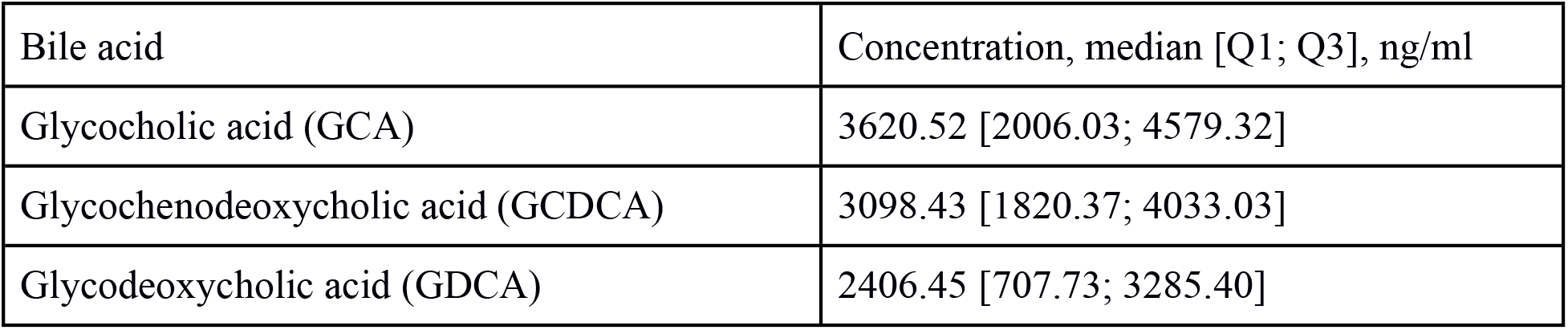
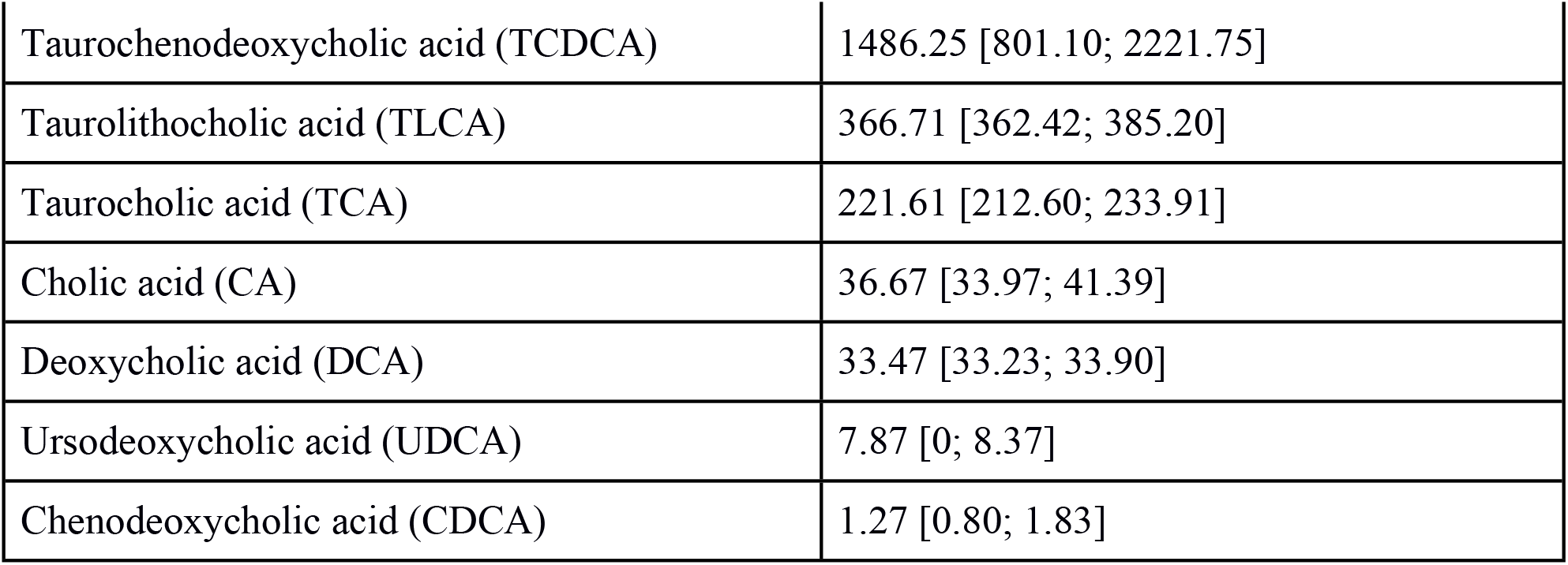
Gallbladder bile acids signature of analyzed bile samples.

### Bile acids and microbial community structure

As it was shown recently, *O. felineus* infection affects beta-diversity of the bile flora [3-4]. Thus, to identify the input of BAs on the microbiota variance we included in the analysis the status of the infection with the liver fluke. Pairwise dissimilarity between communities (beta-diversity) was computed in unweighted UniFrac metric. BAs concentrations were transformed with NMDS to one coordinate vector which was added in adonis model along with *O. felineus* infection status and participants’ gender as covariate (Fig 1). It results in 5.8% of microbial data variance explained with *O. felineus* infection status (p=0.006) and 4.6% of variance explained with BAs (p=0.025). Gender doesn’t provide significant input in microbial community structure (p=0.156).

**Figure 1.**
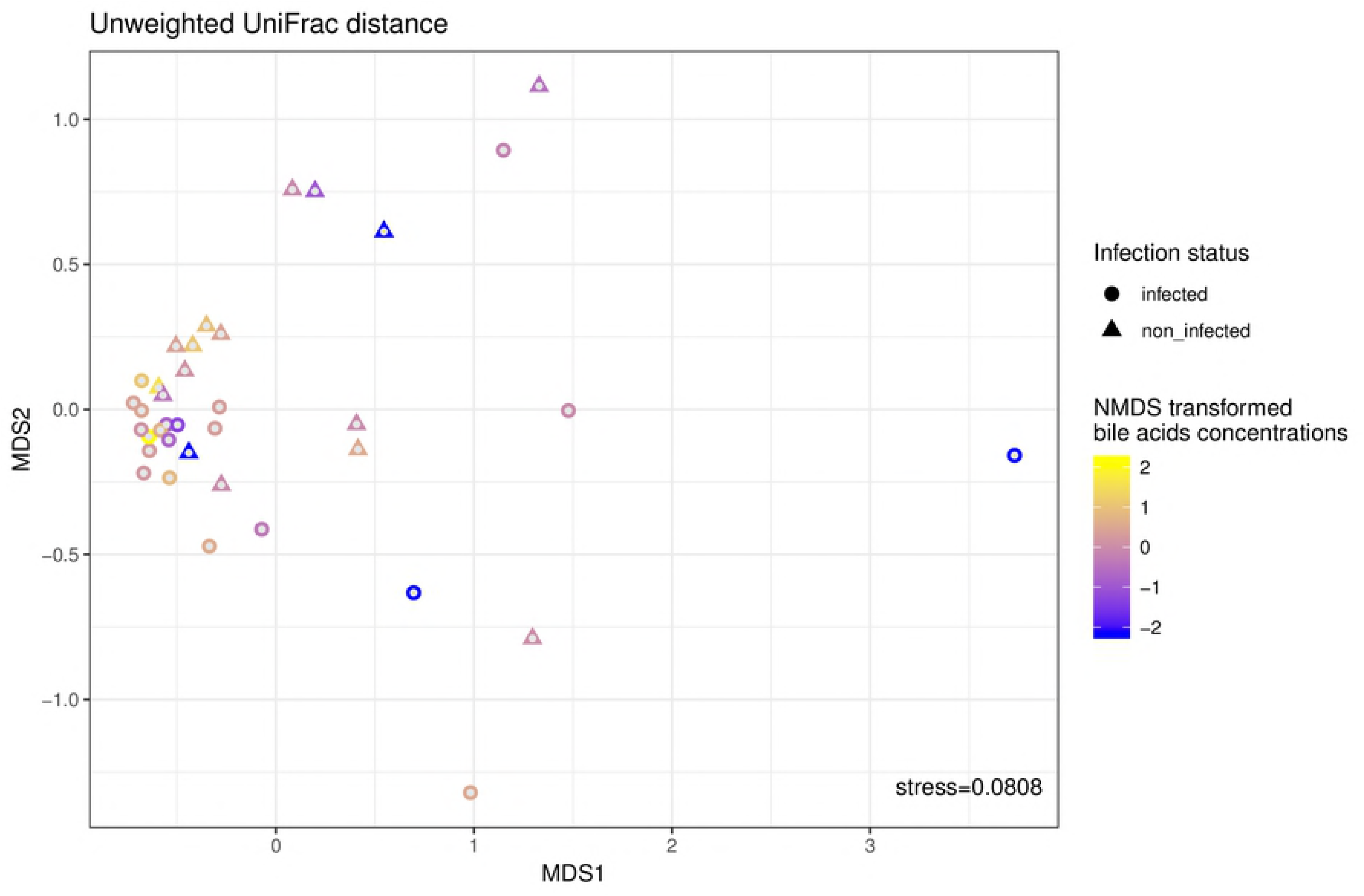
Multidimensional scaling of CSS-corrected OTUs abundances in unweighted UniFrac metric. Circle dots represents *O. felineus* infected samples, triangle dots represents control samples. Color of dots represent value of first principal coordinate of MDS-transformed metabolites, low values showed in blue, high values showed in yellow.

Taxonomic richness (alpha-diversity) of gallbladder microbiota was estimated at a depth of 200 sequences with richness metrics of Chao1, PD whole tree, Shannon, Simpson and a number of observed OTUs. To identify possible associations between microbiome alpha-diversity and BAs levels Spearman correlation was used. It were found significant direct correlations between microbial communities richness measured with Chao1, phylogenetically-driving PD whole tree indices as well as a number of observed OTUs and the levels of taurine-conjugated forms of primary BAs (TCA and TCDCA, Fig 2).

**Figure 2.**
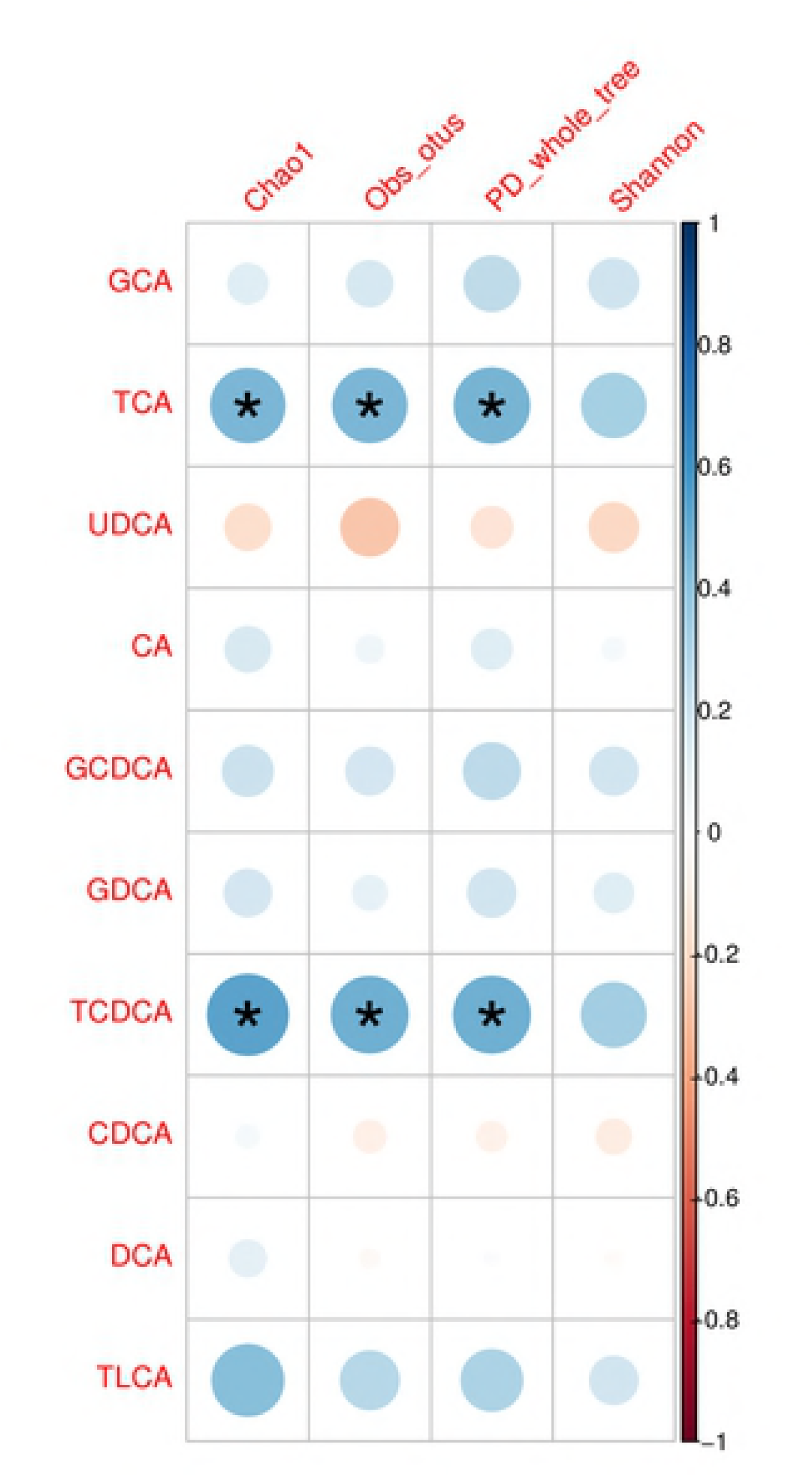
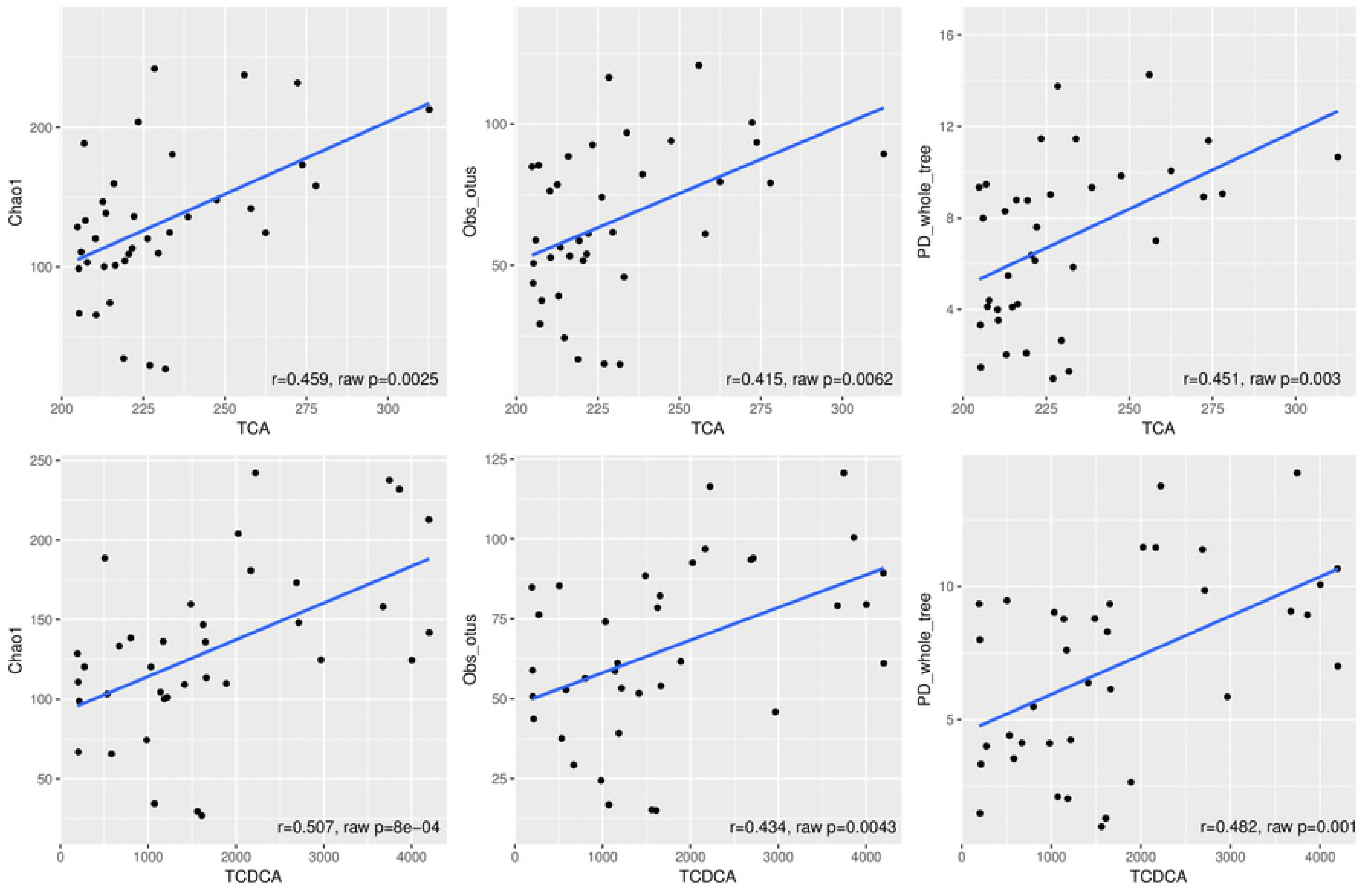
Correlations between alpha-diversity and bile acids concentrations. A) On plot blue circles represents positive correlations, red circles represents negative correlations, significant correlations (FDR-corrected p-value < 0.05) marked with asterisk. B) On plot, blue line represents regression curve for significant associations, gray zone represents standard error. Abbreviations: cholic acid – CA, chenodeoxycholic acid – CDCA, deoxycholic acid – DCA, ursodeoxycholic acid – UDCA, taurocholic acid – TCA, taurochenodeoxycholic acid – TCDCA, taurolithocholic acid – TLCA, glycocholic acid – GCA, glycochenodeoxycholic acid – GCDCA, glycodeoxycholic acid – GDCA, Observed OTUs index – Obs_otus.

### Associations in the system of gallbladder microbiota and bile acids

Thus, considering the results presented in the Figure 2, TCA and TCDCA were used to test the correlations between BA concentration and appearance of bacterial OTUs in bile. For the analysis, bacterial counts were recomputed to presence/absence boolean values. Linear regression revealed associations of bacterial OTUs presence and levels of TCA and TCDCA in gallbladder bile. TCA concentration was directly linked with the presence of Actinomycetales and Bacteroidales orders in gallbladder flora. Chitinophagaceae family, *Lautropia*, *Lutibacterium, Microbacterium* and uncultivated 1-68 genus of [Tissierellaceae] family as well as *Jeotgalicoccus psychrophilus, Prevotella intermedia* and *Haemophilus parainfluenzae* species also were linked with TCA concentration (Table 2, S1 Fig). TCDCA concentration shows positive associations with the presence of OTUs belonging to Chitinophagaceae and Acetobacteraceae families, *Microbacterium, Lutibacterium* and Sphingomonas genera and Prevotella intermedia species (Table 2, S2 Fig).

**Table 2.** associations between BAs levels and microbial OTUs abundances.

| Bile acid | Taxonomy | beta | Standard error | p-value | adj.p-value |
| --- | --- | --- | --- | --- | --- |
| TCA | Actinomycetales | 41.90 | 10.78 | 0.0006 | 0.0286 |
| TCA | *Microbacterium* | 40.69 | 11.52 | 0.0015 | 0.0396 |
| TCA | Bacteroidales | 84.87 | 22.18 | 0.0007 | 0.0286 |
| TCA | *Prevotella intermedia* | 28.81 | 8.37 | 0.0019 | 0.0465 |
| TCA | Chitinophagaceae | 54.63 | 9.89 | 7.51E-006 | 0.0028 |
| TCA | *Jeotgalicoccus psychrophilus* | 47.64 | 12.96 | 0.0010 | 0.0318 |
| TCA | 1-68 of [Tissierellaceae] | 48.09 | 12.82 | 0.0009 | 0.0313 |
| TCA | *Lutibacterium* | 46.67 | 12.15 | 0.0007 | 0.0286 |
| TCA | *Lautropia* | 64.86 | 14.15 | 9.30E-005 | 0.0114 |
| TCA | *Haemophilus parainfluenzae* | 84.87 | 22.18 | 0.0007 | 0.0286 |
| TCDCA | *Microbacterium* | 2114.47 | 550.69 | 0.0007 | 0.0286 |
| TCDCA | *Prevotella intermedia* | 1548.85 | 393.14 | 0.0005 | 0.0286 |
| TCDCA | Chitinophagaceae | 2608.39 | 501.50 | 1.77E-005 | 0.0033 |
| TCDCA | Acetobacteraceae | 2399.28 | 652.76 | 0.0010 | 0.0318 |
| TCDCA | *Lutibacterium* | 2178.51 | 613.07 | 0.0014 | 0.0396 |
| TCDCA | *Sphingomonas* | 1310.20 | 386.61 | 0.0022 | 0.0499 |

In table, beta represents regression line's slope coefficient, SE represents beta's standard error, adj.p-value represents FDR-corrected p-value.

### Bile acids concentrations and *O. felineus* infection

The results of metabolomics analysis of *O. felineus* infection in animal model show that urinary metabolic profiles of the experimental animal change considerably and the urinary BAs are among the main factors explaining the effect [18]. Thus, to investigate a role of the infection status on BAs composition in gallbladder disease we analyzed the measured BAs with respect to the infection status of the patients. There were no significant differences in BAs level (Fig 3), total gallbladder BAs concentration (p=0.24), primary to secondary BAs ratio (p=0.59) between *O. felineus* infected patients and control group.

**Figure 3.**
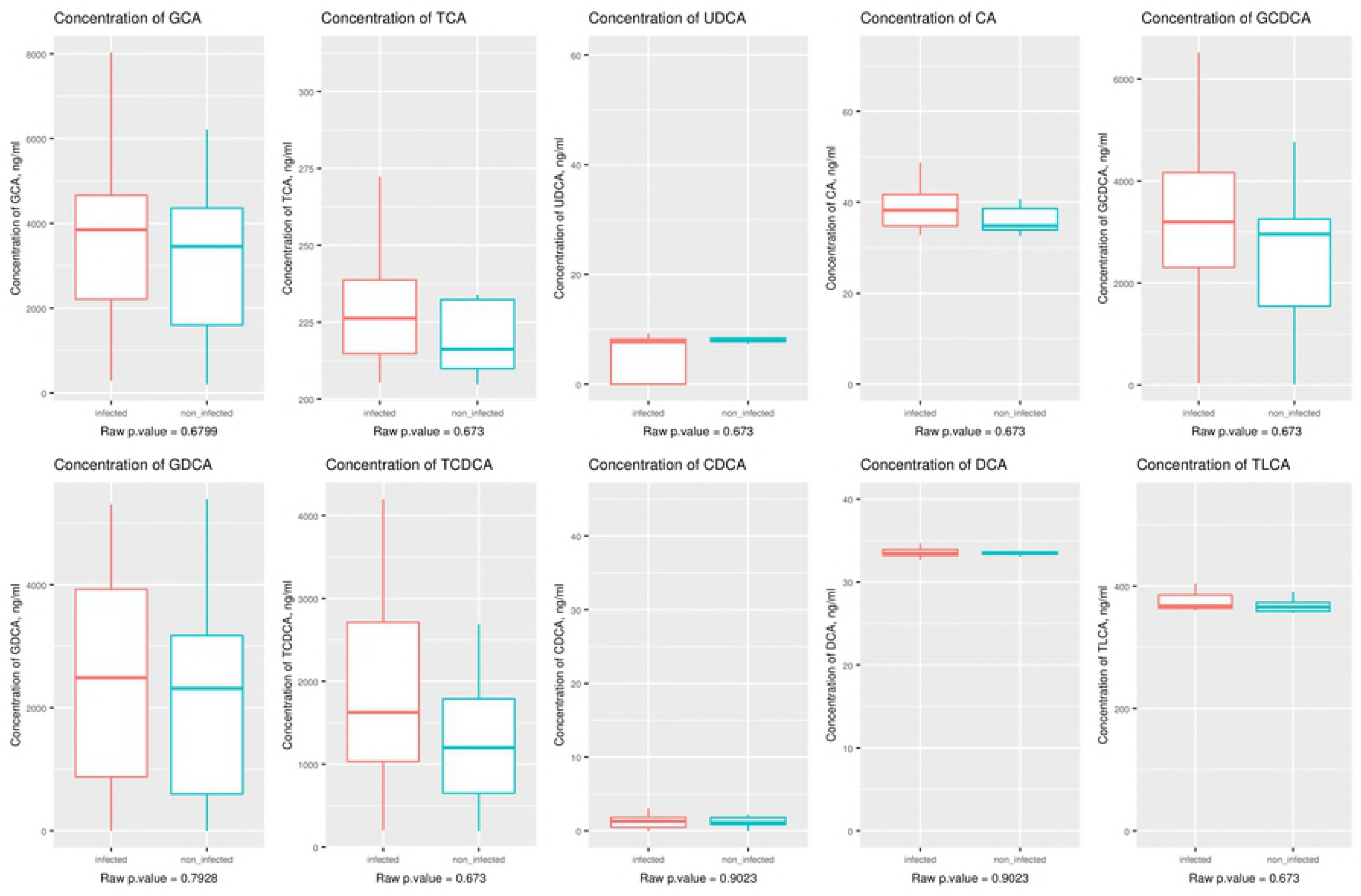
Concentrations of bile acids in gallbladder bile samples. On the plot red boxes represents patients with *O. felineus* infection, blue boxes represents non-infected subjects. Whiskers length represents 1.5 of interquartile range. Abbreviations: cholic acid – CA, chenodeoxy cholic acid – CDCA, deoxycholic acid – DCA, ursodeoxycholic acid – UDCA, taurocholic acid – TCA, taurochenodeoxycholic acid – TCDCA, taurolithocholic acid – TLCA, glycocholic acid – GCA, glycochenodeoxycholic acid – GCDCA, glycodeoxycholic acid – GDCA.

## Discussion

A functional interplay between BAs and microbial composition in gut is a well-documented phenomenon [19]. In bile this phenomenon is far less studied and with this report we describe the interactions between the BAs and microbiota in this complex biological matrix. Collecting the material for this study we could not avoid inclusion of the patients with O. felineus infection, thus it is only logical that we stress a possible influence of the infection on BAs and bile microbiota community. The role of *O. felineus* infection in the bile microbiota composition was proposed and discussed by Saltykova et al., 2016 [4]. We show that *O. felineus* infection has no strong effect on the BAs profile in patients with cholelithiasis. Yet, the infection influences the galbladder microbiota beta-diversity.

Furthermore, we reported significant direct correlations between TCA and TCDCA and the bile microbiota alpha-diversity. TCA concentration was associated with the appearance of species *Jeotgalicoccus psychrophilus, Prevotella intermedia* and *Haemophilus parainfluenzae* in the bile. TCDCA concentration shows positive associations with the presence of OTUs belonging to *Microbacterium, Lutibacterium* and *Sphingomonas* genera and *Prevotella intermedia* species.

In our study we observed correlations between primary BAs and bile bacteria, while fecal microbiota disturbance associated mostly with secondary BAs in feces. Specifically, the analysis of BAs and fecal microbiota in gallstone patients revealed that genus *Oscillospira* was positively correlated with the fraction of secondary BAs, this association attributed to the association of *Oscillospira* and relative fraction of lithocholic acid in the feces [20]. The positive correlation between bacterial taxa and secondary BAs was observed for patients with alcoholic cirrhosis and severe alcoholic hepatitis [21]. It was hypothesized, that microbiota and secondary BAs correlations observed due to the role of the gut flora in the direct or indirect conversion of primary BAs to secondary BAs [20]. Here we observed correlations between primary BAs and bile microbiota that may be related to different mechanisms of the selective force of BAs for bile and gut microbiota.

It is worth of mentioning that TCA and TCDCA associated with bile microbiota diversity and composition are also known as the markers of liver injury and/or liver dysfunction. A recent report of Luo et al indicates a possible role of TCA as a maker of the liver impairment [22]. Several metabolomic studies demonstrated that TCA and TCDCA concentration elevated in serum of liver cirrhotic patients, and positively correlated with Child–Pugh scores [23]. Moreover, *in vitro* experiments indicate that exposure to TCDCA increases expression of the c-myc oncoprotein in WRL-68 cell (hepatocyte like morphology) and downregulates expression CEBPα tumor suppressor protein in HepG2 cells (epithelial morphology) [24]. In mouse model of hepatocellular carcinoma, augmentation of TCDCA intestinal excretion prevented carcinoma development [24].

Most of bacteria associated with TCA and TCDCA concentrations are treated as opportunistic pathogens. *Haemophilus parainfluenzae* is common liver pathogen. It was found more abundant in fecal microbiome of biliary cirrhosis patients [25] and was isolated from bile samples of patients with acute cholecystitis [26] and liver abscesses [27]. *Microbacterium* genus abundance was associated with different inflammatory disorders: otitis externa [28], noma [29], and bacteremia [30]. OTUs belonging to [Tissierellaceae] family were more abundant in ulcerative colitis sites [31]. *Prevotella intermedia* was identified in atopic liver abscess and in case of periodontitis [32].

In conclusion, associations between diversity, taxonomic profile of bile microbiota and bile BAs levels were evidenced in patients with cholelithiasis. Increase of TCDCA and TCA concentration correlates with bile microbiota alpha-diversity and appearance of opportunistic pathogens in bile of patients with cholelithiasis.

## Acknowledgement

This work was supported by the Russian Science Foundation (project No 14-15-00247)

## Supporting information

**Figure S1 – Concentration of TCA depending on selected OTUs presence in gallbladder microbiota.** Blue boxes represents the samples with the presence of OTU; red boxes represents the samples without of OTU. Whiskers length represents 1.5 of interquartile range

**Figure S2 – Concentration of TCDCA depending on selected OTUs presence in gallbladder microbiota. Blue boxes represents the samples with the presence of OTU; red boxes represents the** samples without of OTU. Whiskers length represents 1.5 of interquartile range

**S1 file Bile acid analysis** LC-MS/MS procedures

## List of Abbreviations

BAs: bile acids
LC-MS/MS: liquid chromatography-mass spectrometry and tandem mass spectrometry
TCA: taurocholic acid
TCDCA: taurochenodeoxycholic acid
OTUs: Operational Taxonomic Units
CA: cholic acid
CDCA: chenodeoxycholic acid
DCA: deoxycholic acid
UDCA: ursodeoxycholic acid
TLCA: taurolithocholic acid
GCA: glycocholic acid
GCDCA: glycochenodeoxycholic acid
GDCA: glycodeoxycholic acid
DCA-d4: deoxycholic acid d-4
NMDS: non-metric multidimensional scaling

